# Aberrant Modular Dynamics of Functional Networks in Schizophrenia and Their Relationship with Neurotransmitter and Gene Expression Profiles

**DOI:** 10.1101/2025.01.25.634845

**Authors:** Yi Zhen, Yaqian Yang, Yi Zheng, Zhiming Zheng, Hongwei Zheng, Shaoting Tang

## Abstract

**Introduction:** Numerous studies have emphasized the time-varying modular architecture of functional brain networks and its relevance to cognitive functions in healthy participants. However, how brain modular dynamics change in schizophrenia and how these alterations relate to neurotransmitter and transcriptomic signatures have not been well elucidated.

**Methods:** We harmonized resting-state fMRI data from a multi-site sample including 223 patients and 279 healthy controls and applied the multilayer network method to estimate the regional module switching rate (flexibility) of functional brain connectomes. We examined aberrant flexibility in patients relative to controls and explored its relations to neurotransmitter systems and postmortem gene expression.

**Results:** Compared with controls, patients with schizophrenia had significantly higher flexibility in the somatomotor and right visual regions, and lower flexibility in the left parahippocampal gyrus, right supramarginal gyrus, right frontal-operculum-insula, bilateral precuneus posterior cingulate cortex, and bilateral inferior parietal gyrus. These alterations were associated with multiple neurotransmitter systems and weighted gene transcriptomic profiles. The most relevant genes were preferentially enriched for biological processes of transmembrane transport and brain development, specific cell types, and previously identified schizophrenia-related genes.

**Conclusions:** This study reveals aberrant modular dynamics in schizophrenia and its relations to neurotransmitter systems and schizophrenia-related transcriptomic profiles, providing insights into the understanding of the pathophysiology underlying schizophrenia.

## Introduction

Schizophrenia (SZ) is a severe polygenic mental disorder that affects approximately one percent of the world’s population,^1^ characterized by hallucinations, delusions, disturbances in thinking and behavior, and cognitive deficits.^2,3^ Extensive neuroimaging studies have reported reproducible abnormalities in structural and functional connectivity,^4,5^ supporting the view of SZ as a connectome disconnection syndrome.^6^ These connectivity disruptions in SZ are involved with a diverse set of brain networks (e.g., the somatomotor and default mode networks),^7,8^ suggesting abnormalities in the interaction between distributed regions instead of focal connectome alterations.^9^ Through the lens of brain networks, these studies offer valuable information for recognizing the sophisticated pathophysiology of SZ.

Recently, studies on brain networks of SZ began to focus on the temporal characteristics of functional connectome.^10–12^ The time-varying features of functional connectome are linked to high-order cognitive functions and task performance,^13,14^ holding promise for understanding the cognitive dysfunctions of SZ. Multilayer network modeling is an appealing method for tracking the temporal dynamic reconfiguration of functional networks, which detects the time-varying community structure of functional connectome and characterizes the regional module switching rate, often referred to as flexibility.^15,16^ Previous studies have related flexibility to a wealth of behavioral traits, including positive affect,^17^ personality dimensions,^18^ learning,^15^ divergent creative thinking,^19^ and cognitive performance.^20^ Flexibility is considered to characterize cognitive flexibility,^16,21^ which represents the ability to adjust our thinking and behavior to the environment.^22^ Altered flexibility has been found in various brain disorders, such as autism,^23^ attention deficit hyperactivity disorder,^21^ and multiple sclerosis.^24^ Prior studies have reported higher brain flexibility in patients with SZ than controls during working memory and resting states.^25–26^ Nevertheless, these studies were generally based on a relatively small sample size of patients. A recent study has shown altered flexibility in untreated first-episode SZ,^27^ but the relationship between the alterations of flexibility and gene expression as well as neurotransmitter systems remains unclear.

Neurotransmitter imbalances are thought to play an important role in the pathophysiology of SZ.^28^ Previous studies have reported regional neurotransmitter abnormalities in SZ, such as lower dopamine and glutamate levels in the frontal cortex.^29^ Recent evidence has indicated a link between neurotransmitter systems and brain flexibility. For instance, it has been shown that healthy individuals have significantly higher flexibility after receiving N-methyl-D-aspartate (NMDA) receptor antagonists.^25^ One recent work suggested that dopaminergic withdrawal in Parkinson’s disease increased local flexibility in the frontal, parietal, and insular regions.^30^ However, it is unclear whether the abnormalities of flexibility in SZ are associated with specific neurotransmitter systems, despite the observation that both flexibility and neurotransmitter levels are altered in SZ.

It has been well established that SZ is a highly heritable disease,^31^ with heritability estimates of around 80% in the classic twin and family studies.^32^ Genome-wide association analysis has identified more than 100 risk genetic loci for SZ, promoting our understanding of the intricate etiology of SZ.^33^ Recent studies in imaging transcriptomics have attempted to integrate gene expression data and alterations in brain structures^34^ and functional connectivity^35^ to reveal the mechanisms behind abnormal neuroimaging phenotypes in SZ. However, the relationship between abnormal flexibility in SZ and gene transcription profiles remains unclear. Preliminary evidence implied that healthy individuals with high SZ genetic risks exhibited altered flexibility during working memory.^25,36^ Thus, we speculate that altered resting-state flexibility in SZ is involved with gene expression profiles of previously identified SZ-related risk genes.

Here, we aimed to examine whether resting-state modular dynamics are altered in SZ, and if so, provide further insights into the potential molecular mechanisms of the alterations. Specifically, we harmonized resting-state fMRI data from a relatively large multi-site sample and applied the multilayer network method to estimate brain time-varying modular structures and flexibility. We hypothesized that patients with SZ would exhibit significantly different module dynamics from healthy controls (HC). Then, we explored the potential link between the altered flexibility in SZ and neurotransmitter systems (neurotransmitter receptors\transporters). Finally, we used a partial least squares (PLS) regression to test the association between the alterations of flexibility and gene expression profiles.

## Material and Methods

### Participants and Data acquisition

We used resting-state fMRI data from eight different publicly available datasets.^37–42^ Through stringent image quality control (see below and Supplementary Methods), a total of 502 participants including 223 patients with SZ and 279 healthy controls were selected for subsequent analyses. Informed consent was obtained from all participants and all data collections were approved by the institutional review boards corresponding to data acquisition sites. The demographic and all participants were presented in Supplementary Table S1. More details about the participant diagnosis and MRI data acquisition were shown in Supplementary Methods.

### Data preprocessing

Structural and functional images were preprocessed using fMRIPrep 20.2.3.^43^ In brief, the preprocessing steps of structural images included intensity non-uniformity correction, skull stripping, brain tissue segmentation, cortical surface reconstruction, and spatial normalization to standard spaces. The steps of functional images included head motion correction, slice-timing correction, alignment between functional and structural images, and resampling to standard spaces. The preprocessed functional data were further performed detrending, band-pass temporal filtering, and nuisance regression.^44–46^ The nuisance regression was conducted orthogonally to temporal filtering.^47^ Participants were excluded from this study if met any of the following criteria: (1) poor brain extraction, (2) bad structural image segmentation, (3) inaccurate registration between functional and structural data, and (4) excessive motion in functional images (mean framewise displacement^48^ (FD) > 0.3 mm or maximal FD > 3 mm). Finally, 300 cortical^49^ and 16 subcortical^50^ parcels were used to parcellate functional data to extract regional average time series. For more details on data preprocessing see Supplementary Methods.

### Tracking modular dynamics of brain network

For each participant, a multilayer functional network was constructed using a sliding window method.^51^ A window length of 60 seconds and a shift step of one repetition time were used for the main analysis.^26,52^ Functional connectivity matrices were generated for each window by calculating the Pearson correlation between regional average time series. In alignment with prior studies,^20,53^ we only retained positive values of connectivity matrices in the main analysis. A generalized Louvain community detection algorithm^54^ was applied to track dynamic modules of a multilayer functional network. The optimal community assignment was determined by maximizing the multilayer modularity. The flexibility of each brain node was defined by the percentage of time windows in which node transitions between different network assignments.^15,21^ Consistent with previous studies,^20,21,23^ the topological resolution parameter *γ* and temporal coupling parameter *ω* of the generalized Louvain community detection algorithm were set to *γ* = 1 and *ω* = 1 in the main analysis.

Given the pseudo-randomness of the generalized Louvain algorithm, we repeated the community detection procedure 100 times.^21,23^ The average flexibility over the 100 instances was used for subsequent analysis. The details of multilayer network community identification can be found in Supplementary Methods.

### Case-control comparison

Considering that the fMRI data were collected from different sites, we first corrected for multi-site effects using the NeuroComBat^55^ harmonization method, with age, group, sex, and mean FD as covariates. We then applied linear models to assess the differences in flexibility between patients with SZ and HC at global, network, and nodal levels. Case-control comparisons controlled for mean FD and demographic regressors including age and sex. The t-statistic of case-control differences was further converted into the effect size, Cohen’s d.^56^ Multiple comparison corrections were performed using the FDR method, and the significance threshold was set to corrected p < 0.05.

### Meta-analysis

To explore the functional implication of brain regions with statistically significant case-control differences, we performed a Neurosynth meta-analysis using the Python package NiMARE.^57,58^ The analysis was conducted separately for regions with significant positive SZ-HC differences and regions with significant negative SZ-HC differences. We only retained the top 15 terms relevant to cognitive behaviors or functions.

### Relation between neurotransmitter systems and case-control differences

To discover potential links between neurotransmitter systems and altered flexibility, we examined spatial correlations between SZ-HC differences in flexibility and the densities of neurotransmitter receptors and transporters. A total of 19 different neurotransmitter receptor and transporter density maps were obtained from one recent work.^59^ The statistical significance of spatial correlation was calculated by 10, 000 spin permutation tests.^60,61^ The FDR method was applied to correct for multiple comparisons.

### Transcriptomic neuroimaging association analysis

We capitalized on transcriptomic data from the Allen Human Brain Atlas (AHBA) database^62^ to assess the relationship between SZ-HC differences in flexibility and gene expression profiles. The AHBA database contained microarray expression data from six postmortem human brains. Given that only two donors had gene expression data for the right hemisphere, we restricted the analysis to the left hemisphere. The gene expression data were preprocessed by the abagen toolbox and aligned to the functional parcellation we used,^63,64^ eventually resulting in expression maps of 15633 genes in 158 brain parcels. We used a partial least squares (PLS) regression to examine the association between SZ-HC differences in flexibility and gene transcription profiles. The PLS regression extracted components that were related to SZ-HC differences in flexibility from transcription profiles of 15633 genes. The first PLS component (PLS1), expressed as a weighted sum of gene transcription profiles, was most relevant to SZ-HC differences in flexibility. The statistical significance of the variance explained by PLS1 and the statistical significance of spatial correlations between PLS1 scores and SZ-HC differences were assessed by 10, 000 spin permutation tests,^60,61^ respectively. The variability of PLS1 weight was estimated using 1, 000 bootstrap resamples of 158 brain parcels.^34,65^ The Z score of PLS1 weight for each gene was evaluated as the weight divided by its bootstrap standard deviation. Consistent with previous studies,^65^ we selected genes with Z > 5 (PLS1+ gene list, *p_FDR_* < 0.001, one-sample Z tests) and Z < -5 (PLS1-gene list, *p_FDR_* < 0.001, one-sample Z tests) for subsequent analysis. More details on transcriptomic data preprocessing and the PLS regression can be found in Supplementary Methods.

To understand the functional implications of the selected genes, we performed functional enrichment analysis for the PLS1+ and PLS1- gene lists, respectively, using the online toolbox ToppGene Suite.^66^ The searched annotation categories included Gene Ontology (GO) Biological Process and Disease. The results of enrichment analysis were corrected by the FDR method, and the significance threshold was set to corrected p < 0.05. Next, we examined whether the selected genes were overexpressed in specific cell types. We obtained seven different cell types from a prior study,^67^ including oligodendrocyte precursors, oligodendrocytes, inhibitory neurons, excitatory neurons, microglia, endothelial cells, and astrocytes. We intersected the PLS1+ or PLS1- gene list with the gene list of each cell type to calculate the number of overlapping genes. Statistical significance was evaluated through a null distribution, which was constructed by repeating the aforementioned calculation of the empirical overlap count 10, 000 times on random gene lists.^68^ A random gene list was generated by randomly sampling the same number of genes as the PLS1+ or PLS1- gene lists from all AHBA genes.

We examined the association between altered flexibility and transcriptionally SZ-related dysregulated genes. The transcriptionally dysregulated genes and their differential gene expression data were obtained from two prior case-control studies in postmortem brain tissues.^69,70^ First, we tested whether the PLS1+ and PLS1- gene lists were enriched for SZ-related dysregulated genes using the same procedure as for the cell type analysis. Next, using Spearman correlation analysis,^71^ we assessed the spatial correspondence between PLS1 weights and disease-related differential gene expression in six diseases including SZ, bipolar disorder, major depression disorder, autism, alcohol abuse disorder, and inflammatory bowel disease. Statistical significance was assessed by 10, 000 permutation tests with randomly shuffled differential gene expression.

### Relation between flexibility and clinical data

We calculated Pearson correlations between flexibility and clinical data including antipsychotic doses, and Positive and Negative Syndrome Scale (PANSS) positive and negative scores. Based on the formula from Leucht et al.,^72^ antipsychotic doses for all patients were converted to chlorpromazine (CPZ) equivalents. In addition, utilizing support vector regression and leave-one-out cross-validation,^23,73^ we tested whether flexibility could predict the PANSS positive and negative scores. More details are shown in Supplementary Methods.

### Validation analysis

We validated our findings under multiple methodological considerations, including window length, parameters of a generalized Louvain algorithm, and thresholding windowed functional connectivity. More details can be found in Supplementary Methods.

## Results

Figure 1A displays average nodal flexibility for the SZ and HC groups, which is highly similar to brain flexibility maps described in previous studies.^23^ At the global level, there was no significant SZ-HC difference in average flexibility (Cohen’s d = -0.01, p = 0.906) (Figure 1B). We grouped brain nodes into eight functional networks.^74^ Network-level analysis revealed that patients showed significantly increased flexibility in the somatomotor network relative to HC (Cohen’s d = 0.27, FDR-corrected p = 0.023) (Figure 1C and Supplementary Table S2). Node-level analysis showed that, compared with the HC group, the SZ group exhibited significantly greater flexibility in the right visual and somatomotor regions including the bilateral paracentral lobule, bilateral precentral gyrus, and bilateral postcentral gyrus, but lower flexibility in the left parahippocampal gyrus, right supramarginal gyrus, right frontal-operculum-insula, and several default mode regions including bilateral precuneus posterior cingulate cortex (pCunPCC) and bilateral inferior parietal gyrus (FDR-corrected p < 0.05) (Figure 1D and Supplementary Table S3). Functional decoding analysis based on the NeuroSynth database suggested regions with higher flexibility were mainly associated with motor functions, whereas regions with lower flexibility were primarily related to memory retrieval, autobiographical memory, and episodic memory (Figure 1E and Supplementary Table S4).

**Figure 1.**
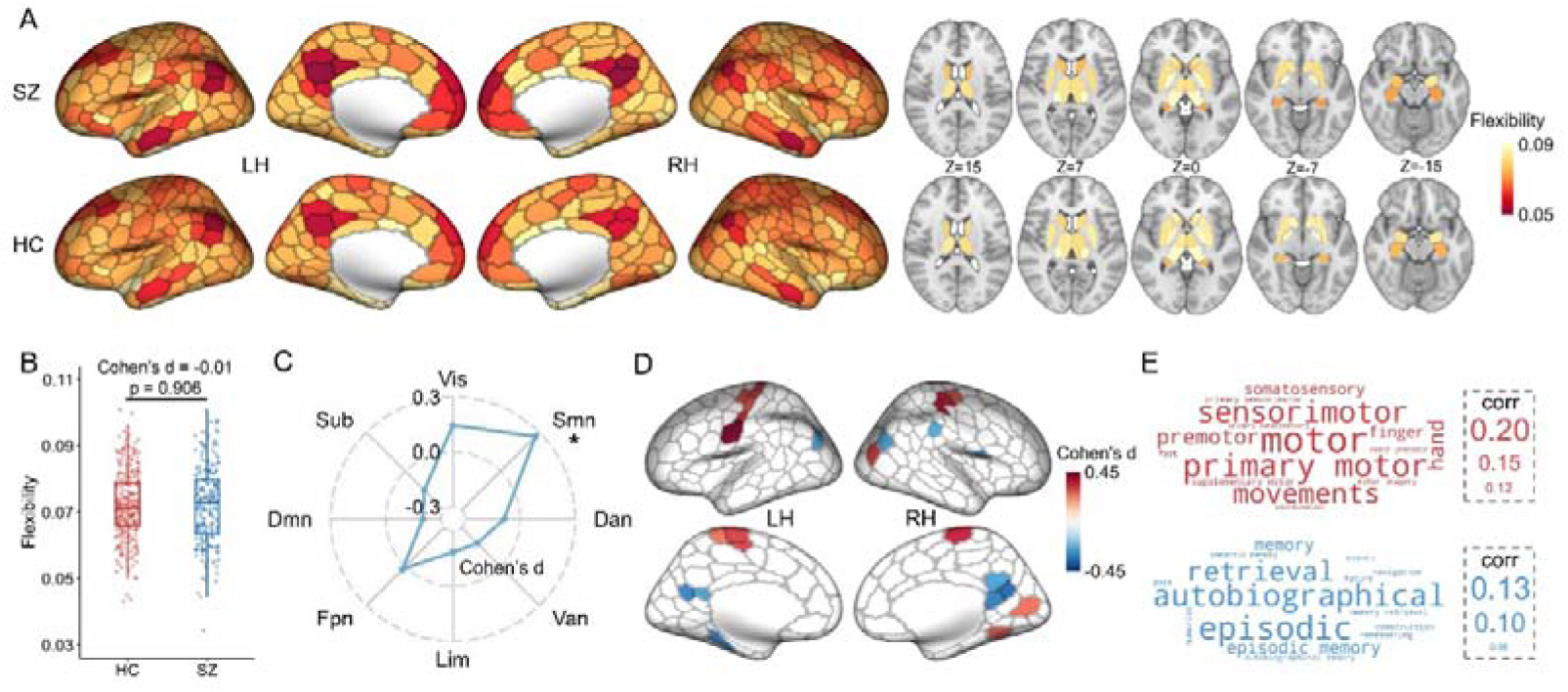
Abnormal modular dynamics in schizophrenia. A. Group-averaged flexibility for patients with schizophrenia (SZ) and healthy controls (HC). B. SZ-HC differences in average flexibility. Boxplots are combined with scatter plots, where each point denotes average flexibility of a participant, the central line of each box indicates the median, the top/bottom edge of each box denotes the 75th/25th percentiles, and the whiskers of each box indicate the upper/lower bounds of 1.5×the interquartile range. C. SZ-HC differences in flexibility across 8 resting-state functional networks (Vis visual network, Smn somatomotor network, Dan dorsal attention network, Van ventral attention network, Lim limbic network, Fpn frontoparietal network, Dmn default mode network, Sub subcortical network). * denotes the statistical significance (FDR corrected p < 0.05). D. significant SZ-HC differences in flexibility at the node level (FDR corrected p < 0.05). E. Cognitive terms associated with regions that exhibit with higher (red) or lower (blue) flexibility in SZ.

Given that neurotransmitter imbalances are implicated in the pathology of SZ, we asked whether SZ-HC differences in flexibility were associated with certain neurotransmitter systems. Figure 2 shows spatial correspondences between the differences in flexibility (Cohen’s d map) and distribution maps of 19 neurotransmitter receptors and transporters. We observed that the Cohen’s d map was significantly positively associated with the density of norepinephrine transporter (NET), and negatively correlated with the densities of several serotonin systems including 5-*HT*_1A_, 5-*HT*_2A_, 5-*HT*_4_, and 5-*HT*_6_, dopamine *D*_1_ and *D*_2_ receptors, cannabinoid *CB*_1_ receptor, acetylcholine *M*_1_ receptor, metabotropic glutamate mGluR5 receptor, and opioid MOR receptor.

**Figure 2.**
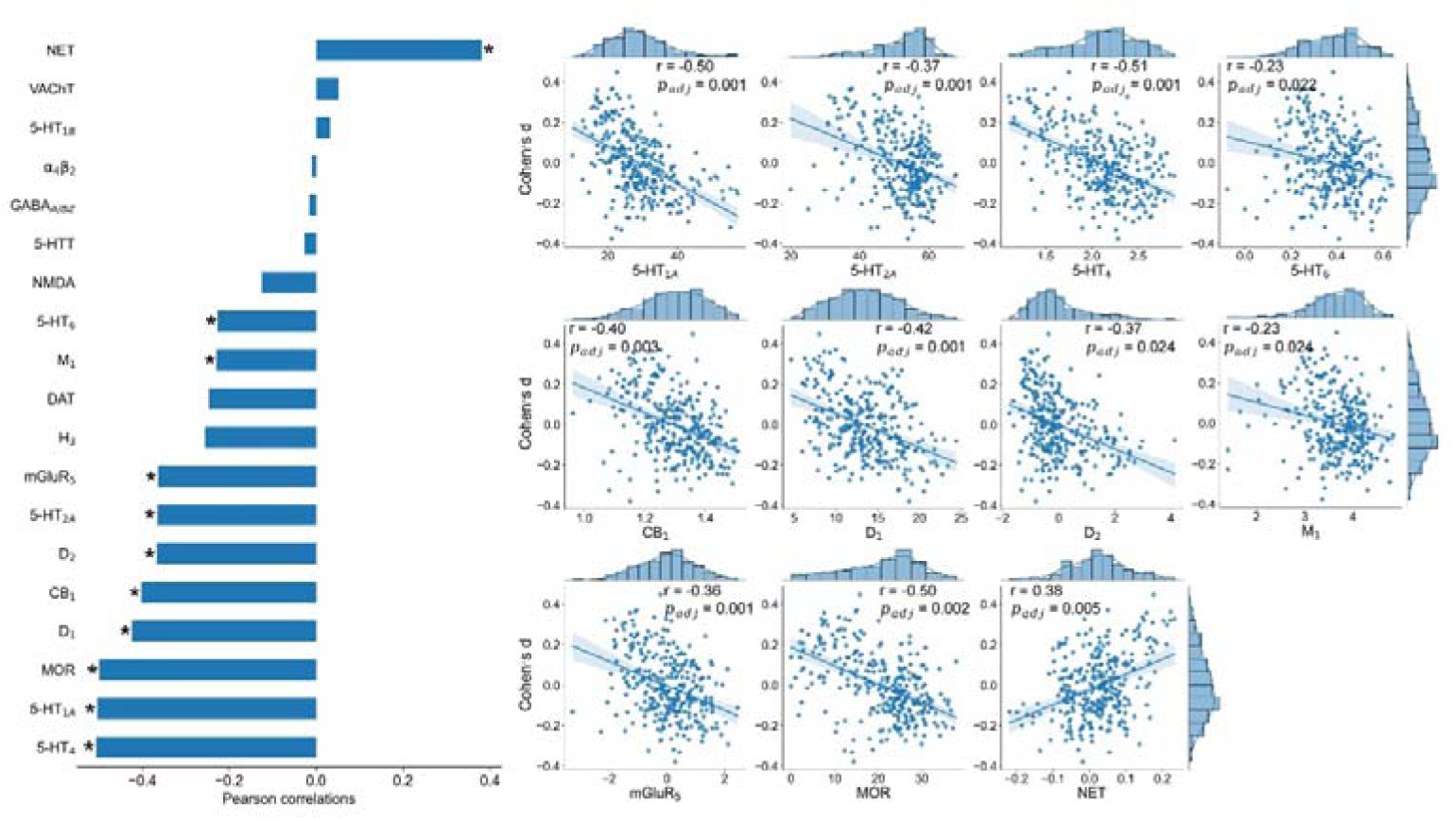
Spatial correlations between SZ-HC differences in flexibility (Cohen’s d map) and the distribution maps of 19 different neurotransmitter receptors and transporters. * denotes statistical significance (FDR corrected p < 0.05).

Schizophrenia is a complex polygenic disorder with diverse genetic variations likely driving the risk of developing schizophrenia. Using postmortem data from the Allen Human Brain Atlas (AHBA) and the partial least squares (PLS) regression, we examined whether the abnormalities of flexibility in SZ were associated with gene expression patterns. We found that the first PLS component (PLS1) accounted for 31.1% of the variance in SZ-HC differences in flexibility (p = 0.0001, 10, 000 spin permutation tests) (Figure 3B). Additionally, the PLS1 weighted gene expression pattern was significantly correlated with SZ-HC differences in flexibility (Pearson’r = 0.56, p < 0.0001, 10, 000 spin permutation tests) (Figure 3C).

**Figure 3.**
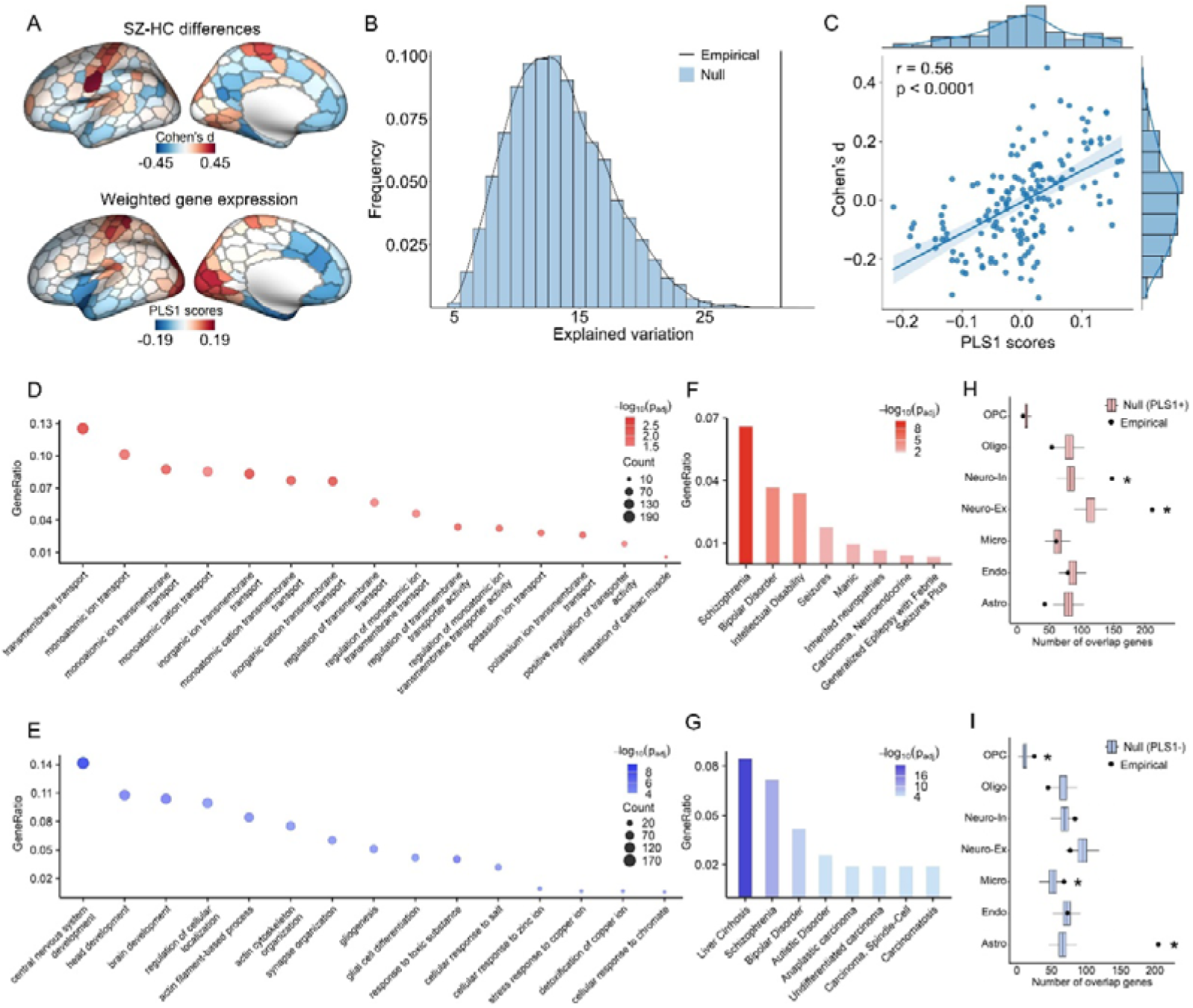
The relationship between the abnormalities of flexibility in SZ and gene expression profiles. A. The graph at the top indicates the SZ-HC cohen’s d map of left hemisphere. The graph at the bottom indicates the weighted gene expression of left hemisphere that derived from the scores corresponding to the first component of the PLS model. B. The significance of variation in SZ-HC differences explained by the first PLS component (p = 0.0001, 10, 000 spin permutation tests). C. Pearson correlation between the PLS1 weighted gene expression pattern and SZ-HC differences in flexibility (p < 0.0001, 10, 000 spin permutation tests). D & E. Biological processes in which the top contributing genes with positive loadings (the PLS1+ genes) (D) and negative loadings (the PLS1- genes) (E) are determined using the Gene Ontology (GO) enrichment analysis. Only the 15 most significant GO terms are shown. The gene ratio (Y-axis) represents the percentage of the number of input genes appearing in the given GO term out of the total number of genes in the category of biological process. F & G. Diseases in which the top contributing genes with positive loadings (the PLS1+ genes) (F) and negative loadings (the PLS1- genes) (G) are identified using the disease enrichment analysis based on the DisGeNET database. Only the 8 most significant terms are shown. The gene ratio (Y-axis) is the percentage of the number of input genes appearing in the given term out of the total number of genes in the category of disease. H & I. Cell types corresponding to the PLS1+ genes (H) and the PLS1- genes (I). * denotes statistical significance (FDR corrected p < 0.05). OPC: oligodendrocyte precursors, Oligo: oligodendrocytes, Neuro-In: inhibitory neurons, Neuro-Ex: excitatory neurons, Micro: microglia cells, Endo: endothelial cells, Astro: astrocytes.

By utilizing the ToppGene Suite, we performed gene list enrichment analyses to assess the biological significance of genes strongly contributing to PLS1. The contributions of genes were defined according to their normalized weights. We extracted 1557 PLS1+ genes (Z>5) and 1290 PLS1- genes (Z<-5). The GO enrichment analysis revealed that the top terms for the PLS1+ gene list mainly related to biological processes involving ion transmembrane transport, and regulation of transmembrane transporter activity (Figure 3D). The PLS1- gene list was primarily enriched for biological processes associated with brain development (Figure 3E). Disease enrichment analyses based on the DisGeNET source^75^ suggested that the PLS1+ genes were predominantly related to schizophrenia, bipolar disorder, and intellectual disability (Figure 3F), while the PLS1- genes were predominantly associated with liver cirrhosis, schizophrenia, and bipolar disorder (Figure 3G). These results informed that the abnormalities of flexibility in SZ were likely to be associated with previously identified SZ-related genes. The detailed results of gene list enrichment analysis can be found in Supplementary Table S5-S6. In addition to their biological functions, we also investigated whether the PLS1+ and PLS1- genes were preferentially enriched in specific cell types. We observed that the PLS1+ genes were more expressed in inhibitory and excitatory neurons (Figure 3H), while the PLS1- genes were more expressed in oligodendrocyte precursors, microglial cells, and astrocytes (Figure 3I).

To further confirm the relationship between aberrant flexibility in SZ and SZ-related genes, we additionally performed two different analyses based on transcriptional data of postmortem brain tissue from two case-control studies of SZ. First, we tested whether the PLS1+ and PLS1- genes were overexpressed in SZ-related dysregulated genes. We found that the PLS1+ genes were significantly enriched for SZ-related downregulated genes reported by Fromer et al.^70^ and Gandal et al.,^69^ and that PLS1- genes were significantly enriched for SZ-related upregulated genes reported by Fromer et al.^70^ and Gandal et al.^69^ as well as SZ-related downregulated genes reported by Fromer et al.^70^ (Figure 4A). Second, we examined spatial associations between PLS1 weights and differential gene expression (DGE) patterns in postmortem brain tissue. The PLS1 weights of PLS1+ and PLS1- genes were significantly negatively correlated with the DGE patterns of SZ-related significantly dysregulated genes reported by Fromer et al.^70^ and Gandal et al.^69^ Although significant associations were observed between the PLS1 weights and the DGE patterns for other disorders (bipolar disorder and autism), the strongest magnitude of associations was found for SZ (Figure 4B). Additionally, we retained all differential gene expressions and PLS1 weights for all genes and assessed their spatial correlations. The results were largely in agreement with those from analyses using only the PLS1+ genes, the PLS1- genes, and significant dysregulated genes (see Supplementary Figure S1). In sum, these findings highlighted a potential link between altered flexibility in SZ and SZ-related genes, consistent with the results of disease enrichment analysis.

**Figure 4.**
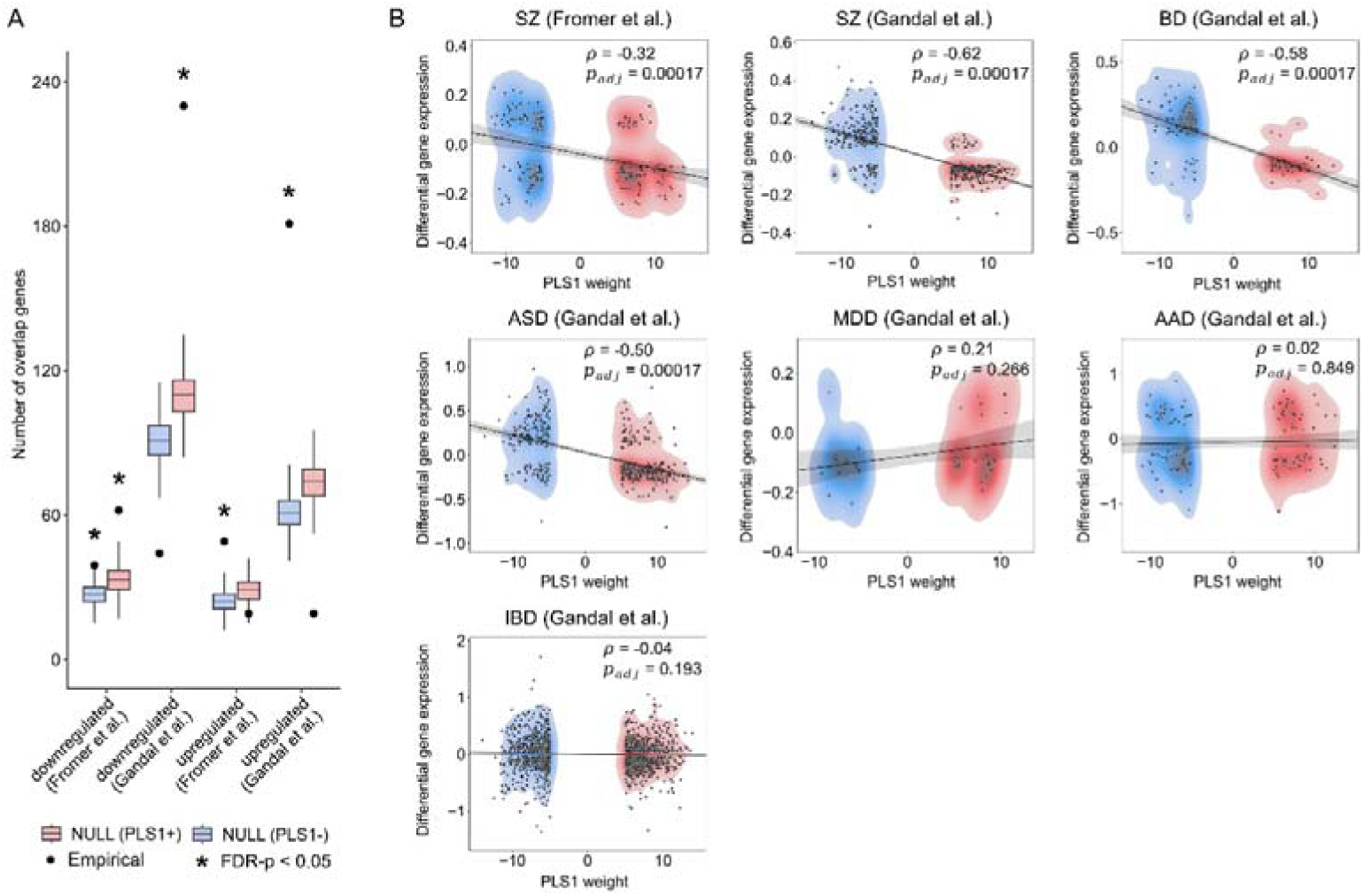
The relationship between altered flexibility in SZ and SZ-related dysregulated genes. A. The number of overlapping genes between PLS1 genes (PLS1+ and PLS1- genes) and significant SZ-related dysregulated genes reported by Gandal et al. and Fromer er al. B. Spearman correlations between the PLS1 weights (PLS1+ and PLS1- gene weights) and the differential gene expression of significant disease-related dysregulated genes reported by Gandal et al. and Fromer er al. SZ: schizophrenia, BD: bipolar disorder, ASD: autism spectrum disorder, MDD: major depression disorder, AAD: alcohol abuse disorder, IBD: inflammatory bowel disease.

Finally, we examined the association between flexibility and clinical measures. There were no significant relations between CPZ equivalent doses and global average flexibility, as well as between CPZ equivalent doses and nodal flexibility. Neither PANSS positive scores nor PANSS negative scores were detected to be significantly correlated with flexibility for global and nodal levels. Utilizing support vector regression and leave-one-out cross-validation, we found that nodal flexibility provided a weak but statistically significant prediction for PANSS positive scores (Pearson’r = 0.20, FDR-corrected p = 0.012), but not for PANSS negative scores (Pearson’r = 0.07, FDR-corrected p = 0.185) (see Supplementary Figure S2).

The main results were largely consistent under multiple confounding factors including sliding window lengths, different parameter settings of the community detection algorithm, and thresholding window functional connectivity (see Supplementary Figure S3-S8).

## Discussion

In the current study, we demonstrated altered time-varying modular reconfigurations in patients with SZ, with significantly increased flexibility in the somatomotor and right visual regions, as well as reduced flexibility in left parahippocampal gyrus, right supramarginal gyrus, right frontal-operculum-insula, bilateral precuneus posterior cingulate cortex, and bilateral inferior parietal gyrus. These abnormalities were related to multiple neurotransmitter systems, including serotonin, dopamine, norepinephrine, cannabinoid, acetylcholine, glutamate, and opioid. Transcriptomic association analyses suggested that aberrant flexibility related to weighted gene expression patterns, with strongly contributing genes enriched for the biological processes of transmembrane transport and brain development, specific cell types (e.g., oligodendrocyte precursors, microglia cells, astrocytes, and inhibitory and excitatory neurons), and previously identified SZ-related genes. Altogether, these findings provide evidence of aberrant modular dynamics in SZ and reveal relations of these abnormalities to neurotransmitter systems and genetic factors, advancing our understanding of the pathophysiology underlying SZ.

Prior studies have consistently reported abnormal static functional connectivity in SZ in the somatomotor and default mode regions.^8,76^ Moving beyond the static methods, our work investigated SZ-related alterations in dynamic modular reconfigurations and demonstrated aberrant flexibility in the primary sensory cortex (e.g., somatomotor and right visual regions) and high-order association cortex (e.g., inferior parietal gyrus and pCunPCC). These findings indicate that, in addition to previously reported impaired functional interactions, aberrant temporal switching of network modular architecture is a significant feature of SZ. The observed widespread abnormalities promote the emerging consensus that neuropathology of SZ involves disruption of multi-network systems beyond focal alterations.^7,9^ In addition, the multilayer network model used here can characterize node-level dynamic configurations, thus extending the existing research that commonly focuses on whole-brain state transients via sliding windows and clustering algorithms.^51,77^ Compared with these findings that SZ significantly differs from HC in dwell times of multiple whole-brain states,^10,78^ our work characterizes spatially relatively refined changes in brain dynamics in SZ. Through functional decoding analyses, we found that regions with higher flexibility were mainly related to motor functions whereas regions with lower flexibility were primarily related to memory functions, implying that the directions of altered dynamic configurations might correspond to distinct clinical symptoms of SZ. However, due to the lack of detailed symptom information in patients, further work is warranted to examine direct associations between altered flexibility and distinct symptoms.

In particular, network-level and node-level analyses suggested that regions with the largest effect size and most statistical significance for case-control differences in flexibility are located in the somatomotor network. Prior imaging studies have reported that patients with SZ have the strongest connectivity disturbances in the somatomotor network, highlighting that targeting the somatomotor system potentially contributes to the neuromodulation treatment of SZ and the understanding of its pathology.^79,80^ Transdiagnostic studies have suggested that the somatomotor system serves as a transdiagnostic hub that relates to cognitive dysfunction, general psychopathology, and impulsivity.^81^ In alignment with these findings, our results also emphasize the essential role of the somatomotor system in SZ neuropathology.

Recent studies using the multilayer network model have highlighted that patients with SZ tend to have higher flexibility during working memory and rest compared to HC.^25–27^ By harmonizing resting-state fMRI data from a relatively large multi-site sample, we extended previous discoveries and revealed bidirectional changes of brain flexibility in schizophrenia. Specifically, we found that SZ exhibited not only increased flexibility primarily in the somatomotor network, which was consistent with recent findings in untreated first-episode SZ,^27^ but also decreased flexibility in some areas mainly involved in the default mode network, implying that SZ might exhibit more nuanced and complex alterations in flexibility than previously reported. As a result, we detected no significant case-control differences rather than previously reported increases in whole-brain average flexibility.^25,27^ Moreover, we detected no significant differences in the subcortical and frontoparietal control networks, whereas one prior study reported increased flexibility in these networks.^26^ These inconsistencies may be partly due to differences in sample size, as small sample sizes potentially limit statistical power and sensitivity. Our analyses were conducted in a relatively large sample including 502 participants, in contrast with previous studies with samples of no more than 250 participants. Patient’s treatment status, duration of illness, and fMRI data acquired under different tasks may also contribute to these discrepancies.^25,27^

Our results revealed a significant spatial concordance between case-control differences in flexibility and the densities of several neurotransmitter systems including serotonin, dopamine, norepinephrine, acetylcholine, glutamate, cannabinoid, and opioid. These neurotransmitter receptors and transporters have been previously reported to be involved with the pathology of SZ.^82–87^ For example, the 5-*HT*_1A_ receptor has been implicated in the regulation of human emotion and cognitive,^88^ and has been reported to show a significant increase in the prefrontal cortex of individuals with SZ compared to healthy controls in post-mortem studies.^89^

The mu-opioid receptor (MOR) is related to reward processing and hedonic function and shows a significantly decreased availability in the striatum of individuals with SZ.^87^ Recent studies have suggested that brain network dynamics are regulated by neurotransmitter receptors involving dopamine, glutamate, and 5-*HT2a*.^25,90,91^ We speculate that neurotransmitter imbalances potentially contribute to abnormal dynamic modular configurations of brain networks in SZ.

By leveraging transcriptome data from the AHBA database, we identified a strong link between case-control differences in flexibility and weighted gene expression. Our findings on the genetic underpinning of abnormal modular dynamics have three implications. First, the most correlated genes with positive weights were primarily enriched for transmembrane transport (that is, the transport of a solute across a lipid bilayer (GO:0055085)). Prior findings at the molecular level have provided evidence that patients with SZ have aberrant mitochondrial channel protein levels^92^ and disrupted glutamate transporter protein complex,^93^ potentially impeding normal solute transport across membranes. The protein-protein interaction (PPI) network analysis has shown that SZ risk loci are related to ion transmembrane transport^94^ and regulation of ion transmembrane transport.^95^ Combining our findings with the aforementioned studies, we posit that cortical regions with altered flexibility in SZ might be associated with abnormal biological processes of transmembrane transport. The most correlated genes with negative weights are mainly enriched for the terms related to brain development, suggesting that abnormal flexibility might correlate with the neurodevelopmental process of SZ. In addition, our findings are potentially in support of the developmental hypothesis of SZ pathogenesis.^96^ Second, by assigning the most correlated genes to specific cell classes, our results indicate that SZ-related alterations in flexibility might be linked to the alterations of specific cell types including oligodendrocyte precursors, microglia cells, and astrocytes, inhibitory and excitatory neurons. It has been previously reported that these cell types are involved in the pathophysiology of SZ.^97–101^ Third, through three different analyses, our results provide robust evidence that altered flexibility in SZ is associated with previously reported SZ-related risk genes. This association was indirectly implied by prior literature under the working memory paradigm. Braun et.al reported that unaffected first-degree relatives of SZ showed significantly increased flexibility compared with HC during working memory.^25^ Dimitriadis et al.^36^ showed that participants with high SZ-related genetic risk had reduced flexibility during working memory. In this work, we provide the first direct evidence of the link between SZ-related genes and abnormal flexibility in SZ during the resting state.

There are several limitations to our findings that need to be considered. First, the large sample we used pooled from multiple sites with different acquisition parameters. Although we utilized the NeuroCombat^55^ method to correct site effects, future studies should use a large sample with the same collection parameters to validate our findings. Second, due to the dearth of detailed information on medication use and duration of illness in many patients, we did not account for these factors in the case-control comparisons. Third, we used neurotransmitter distribution maps and gene expression profiles obtained from participants without SZ. Future studies should integrate neurotransmitter and gene expression data from patients to further validate our findings.

## Conclusions

By harmonizing resting-state fMRI data from a multi-site sample, this study suggests abnormal dynamic modular reconfigurations in SZ, with significantly higher flexibility in the somatomotor and right visual regions, as well as lower flexibility primarily in the default mode regions. The spatial pattern of altered flexibility correlates with the distributions of multiple neurotransmitter systems. Transcriptomic association analyses inform that these alterations are associated with gene transcriptomic profiles preferentially enriched for biological processes of transmembrane transport and brain development, and previously identified SZ-related genes. In sum, this study suggests that schizophrenia is related to abnormal dynamic modular reconfiguration of resting-state functional networks, and reveals that schizophrenia-relevant genes and neurotransmitter systems are associated with these alterations.

## DATA AVAILABILITY STATEMENT

The COBRE data was downloaded from the COllaborative Informatics and Neuroimaging Suite Data Exchange tool (COINS; http://coins.mrn.org/dx) and data collection was performed at the Mind Research Network, and funded by a Center of Biomedical Research Excellence (COBRE) grant 5P20RR021938/P20GM103472 from the NIH to Dr. Vince Calhoun. The BrainGluSchi Data was downloaded from the COllaborative Informatics and Neuroimaging Suite Data Exchange tool (COINS;http://coins.mrn.org/dx) and data collection was funded by NIMH R01MH084898-01A1. This data was obtained from the OpenfMRI database (https://openfmri.org/dataset/ds000030/). Its accession number is ds000030. The SRPBS data were obtained from the DecNef Project Brain Data Repository (https://bicr-resource.atr.jp/srpbsopen/), collected as part of the Japanese Strategic Research Program for the Promotion of Brain Science (SRPBS) supported by the Japanese Advanced Research and Development Programs for Medical Innovation (AMED).

## Acknowledgments

This work is supported by National Science and Technology Major Project (2022ZD0116800), Program of National Natural Science Foundation of China (12425114, 62141605, 12201026, 12301305), the Fundamental Research Funds for the Central Universities, and Beijing Natural Science Foundation (Z230001).

