## Supplementary Materials for "Aberrant Modular Dynamics of Functional Networks in Schizophrenia and Their Relationship with Neurotransmitter and Gene Expression Profiles"

**Supplementary methods**

**Participants and Data acquisition**

COBRE dataset^1^: This open-access dataset was collected at the Mind Research Network. This collection was approved by the institutional review board (IRB) of the University of New Mexico (UNM) and all subjects provided written informed consent. Each participant was diagnosed by the Structured Clinical Interview for DSM-IV Axis I Disorders^2^. The symptom ratings of patients were assessed through the Positive and Negative Syndrome Scale (PANSS)^3^. The olanzapine equivalents were used to estimate the doses of antipsychotic medications. All imaging data were acquired on a Siemens 3 T TIM Trio scanner, using a 12-channel head coil. The T1-weighted images were acquired with a multi-echo MPRAGE sequence (TE = 1.64, 3.5, 5.36, 7.22, 9.08ms, TR = 2.53s, TI = 1.2s, flip angle = 7^o^, slice thickness = 1mm, FOV = 256mm, resolution = 256$\times$256, voxel size = 1.0$\times$1.0$\times$1.0mm). The fMRI data were acquired with a gradient-echo echoplanar pulse sequence (TE = 29ms, TR = 2s, flip angle = 75^o^, FOV = 240mm, 33 slices, matrix size = 64$\times$64, voxel size = 3.75$\times$3.75$\times$4.55mm).

BrainGluSchi dataset^4^: This study was approved by the UNM Institutional Review Board and subjects gave written informed consent. Participants were diagnosed by the DSM-IV-TR^5^. Imaging data were collected on a 3 Tesla scanner (VB-17; 12 channel head-coil). The T1-weighted images were acquired with 3D-MPRAGE (TE = 3.87ms, TR = 1.5s, TI = 700ms, flip angle = 10^o^, slice thickness = 1mm, FOV = 256$\times$256mm). The fMRI data were collected using the following parameters: TE = 29ms, TR = 2s, flip angle = 75^o^, FOV = 240mm, 33 slices, voxel size = 3.8$\times$3.8$\times$3.5mm.

UCLA-CNP dataset^6^: Subjects gave written informed consent in accordance with procedures approved by the Institutional Review Boards at UCLA and the Los Angeles County Department of Mental Health. Participants were diagnosed using the Structured Clinical Interview for DSM-IV^5^. The MRI data were collected on one of two 3T Siemens Trio scanners. The T1-weighted MPRAGE images were acquired with the following parameters: TR = 1.9s, TE = 2.26ms, FOV = 250mm, matrix = 256$\times$256, 176 slices, slice thickness = 1mm. The fMRI data were collected with a T2^*^-weighted echoplanar imaging (EPI) sequence (TR = 2s, TE = 30ms, matrix = 64$\times$64, flip angle = 90^o^, slice thickness = 4mm, 34 slices, FOV = 192mm).

SRPBS database^7^ (SWA, UTO, KUT, KTT): All participants provided written informed consent. The recruitment procedures and experimental protocols for the SWA\UTO\KUT\KTT dataset were approved by the institutional review boards of the Showa University\University of Tokyo Faculty of Medicine\Kyoto University\Kyoto University. Subjects were diagnosed using the DSM-IV-TR^5^ or DSM-5^8^. For the SWA dataset, the imaging data were collected using a 3T Siemens Verio scanner. The T1-weighted images were acquired with the following parameters: TR = 2.3s, TE = 2.98ms, TI = 900ms, flip angle = 9o, FOV = 256mm, matrix = 256$\times$256, voxel size = 1$\times$1$\times$1mm; the fMRI data were collected with the following parameters: TR = 2.5s, TE = 30ms, flip angle = 80^o^, FOV = 212mm, matrix = 64$\times$64, 40 slices, in-plane resolution = 3.3$\times$3.3mm, slice thickness = 3.2mm. For the UTO dataset, the imaging data were collected using a 3T GE MR750w scanner. The T1-weighted images were acquired with the following parameters: TR = 7.7ms, TE = 3.1ms, TI = 400ms, flip angle = 8^o^, FOV = 240mm, matrix = 256$\times$256, voxel size = 1$\times$1$\times$1.2mm; the fMRI data were collected with the following parameters: TR = 2.5s, TE = 30ms, flip angle = 80^o^, FOV = 212mm, matrix = 64$\times$64, 40 slices, in-plane resolution = 3.3$\times$3.3mm, slice thickness = 3.2mm. For the KUT dataset, the imaging data were collected using a 3T Siemens TimTrio scanner. The T1-weighted images were acquired with the following parameters: TR = 2s, TE = 3.4ms, TI = 990ms, flip angle = 8^o^, FOV = 225$\times$240mm, matrix = 240$\times$256, voxel size = 0.9375$\times$0.9375$\times$1.0mm; the fMRI data were collected with the following parameters: TR = 2.5s, TE = 30ms, flip angle = 80^o^, FOV = 212mm, matrix = 64$\times$64, 40 slices, in-plane resolution = 3.3125$\times$3.3125mm, slice thickness = 3.2mm. For the KTT dataset, the imaging data were collected using a 3T Siemens Trio scanner. The T1-weighted images were acquired with the following parameters: TR = 2s, TE = 4.38ms, TI = 990ms, flip angle = 8^o^, FOV = 225$\times$240mm, matrix = 240$\times$256, voxel size = 0.9375$\times$0.9375$\times$1.0mm; the fMRI data were collected with the following parameters: TR = 2s, TE = 30ms, flip angle = 90^o^, FOV = 256$\times$192mm, matrix = 64$\times$48, 30 slices, in-plane resolution = 4$\times$4mm, slice thickness = 4mm.

The Neural Basis of Visual Shape Completion in Schizophrenia and Bipolar Disorder dataset^9-10^: This dataset was downloaded from the openneuro^11^ platform (ds005073 and ds003404). All participants provided written informed consent. The study was approved by the Rutgers University Institutional Review Board. All participants were diagnosed using the DSM-5^8^. The imaging data were collected on a Siemens Tim Trio scanner. The T1-weighted imaging data were collected using the following parameters: TR = 2.4s, TE = 2.31ms, TI = 1s, slice thickness = 0.8mm, flip angle = 8^o^; the multiband EPI data were collected using a 32-channel head coil with the following parameters: TR = 785ms, TE = 34.8ms, flip angle = 55^o^, FOV = 211mm, 60 slices, 2.4mm isotropic voxels, multiband acceleration factor = 6.

Due to the extreme imbalance between the number of healthy controls (HC) and participants with schizophrenia (SZ) in five datasets (UCLA-CNP, SWA, UTO, KUT, and KTT), we performed the optimal group matching using the R package MatchIt^36^.

**Data preprocessing**

Results included in this manuscript come from preprocessing performed using fMRIPrep 20.2.3^12-13^, which is based on Nipype 1.6.1^14-15^.

**Anatomical data preprocessing**

A total of 1 T1-weighted (T1w) images were found within the input BIDS dataset. The T1-weighted (T1w) image was corrected for intensity non-uniformity (INU) with N4BiasFieldCorrection^16^, distributed with ANTs 2.3.3^17^, and used as T1w-reference throughout the workflow. The T1w-reference was then skull-stripped with a Nipype implementation of the antsBrainExtraction.sh workflow (from ANTs), using OASIS30ANTs as target template. Brain tissue segmentation of cerebrospinal fluid (CSF), white-matter (WM) and gray-matter (GM) was performed on the brain-extracted T1w using fast^18^ (FSL 5.0.9). Brain surfaces were reconstructed using recon-all^19^ (FreeSurfer 6.0.1), and the brain mask estimated previously was refined with a custom variation of the method to reconcile ANTs-derived and FreeSurfer-derived segmentations of the cortical gray-matter of Mindboggle^20^. Volume-based spatial normalization to two standard spaces (MNI152NLin6Asym, MNI152NLin2009cAsym) was performed through nonlinear registration with antsRegistration (ANTs 2.3.3), using brain-extracted versions of both T1w reference and the T1w template. The following templates were selected for spatial normalization: FSL’s MNI ICBM 152 non-linear 6th Generation Asymmetric Average Brain Stereotaxic Registration Model^21^ [TemplateFlow ID: MNI152NLin6Asym], ICBM 152 Nonlinear Asymmetrical template version 2009c^22^ [TemplateFlow ID: MNI152NLin2009cAsym],

**Functional data preprocessing**

For each of the 1 BOLD run found per subject (across all tasks and sessions), the following preprocessing was performed. First, a reference volume and its skull-stripped version were generated using a custom methodology of fMRIPrep. Susceptibility distortion correction (SDC) was omitted. The BOLD reference was then co-registered to the T1w reference using bbregister (FreeSurfer) which implements boundary-based registration^23^. Co-registration was configured with six degrees of freedom. Head-motion parameters with respect to the BOLD reference (transformation matrices, and six corresponding rotation and translation parameters) are estimated before any spatiotemporal filtering using mcflirt^24^ (FSL 5.0.9). BOLD runs were slice-time corrected using 3dTshift from AFNI 20160207^25^. The BOLD time-series were resampled onto the following surfaces (FreeSurfer reconstruction nomenclature): fsaverage5, fsaverage. The BOLD time-series (including slice-timing correction when applied) were resampled onto their original, native space by applying the transforms to correct for head-motion. These resampled BOLD time-series will be referred to as preprocessed BOLD in original space, or just preprocessed BOLD. The BOLD time-series were resampled into standard space, generating a preprocessed BOLD run in MNI152NLin6Asym space. First, a reference volume and its skull-stripped version were generated using a custom methodology of fMRIPrep. Grayordinates files^26^ containing 91k samples were also generated using the highest-resolution fsaverage as intermediate standardized surface space. Several confounding time-series were calculated based on the preprocessed BOLD: framewise displacement (FD), DVARS and three region-wise global signals. FD was computed using two formulations following Power^27^ (absolute sum of relative motions) and Jenkinson^24^ (relative root mean square displacement between affines). FD and DVARS are calculated for each functional run, both using their implementations in Nipype (following the definitions by Power et al.). The three global signals are extracted within the CSF, the WM, and the whole-brain masks. Additionally, a set of physiological regressors were extracted to allow for component-based noise correction^28^ (CompCor). Principal components are estimated after high-pass filtering the preprocessed BOLD time-series (using a discrete cosine filter with 128s cut-off) for the two CompCor variants: temporal (tCompCor) and anatomical (aCompCor). tCompCor components are then calculated from the top 2% variable voxels within the brain mask. For aCompCor, three probabilistic masks (CSF, WM and combined CSF+WM) are generated in anatomical space. The implementation differs from that of Behzadi et al. in that instead of eroding the masks by 2 pixels on BOLD space, the aCompCor masks are subtracted a mask of pixels that likely contain a volume fraction of GM. This mask is obtained by dilating a GM mask extracted from the FreeSurfer’s aseg segmentation, and it ensures components are not extracted from voxels containing a minimal fraction of GM. Finally, these masks are resampled into BOLD space and binarized by thresholding at 0.99 (as in the original implementation). Components are also calculated separately within the WM and CSF masks. For each CompCor decomposition, the k components with the largest singular values are retained, such that the retained components’ time series are sufficient to explain 50 percent of variance across the nuisance mask (CSF, WM, combined, or temporal). The remaining components are dropped from consideration. The head-motion estimates calculated in the correction step were also placed within the corresponding confounds file. The confound time series derived from head motion estimates and global signals were expanded with the inclusion of temporal derivatives and quadratic terms for each^29^. Frames that exceeded a threshold of 0.5 mm FD or 1.5 standardised DVARS were annotated as motion outliers. All resamplings can be performed with a single interpolation step by composing all the pertinent transformations (i.e. head-motion transform matrices, susceptibility distortion correction when available, and co-registrations to anatomical and output spaces). Gridded (volumetric) resamplings were performed using antsApplyTransforms (ANTs), configured with Lanczos interpolation to minimize the smoothing effects of other kernels^30^. Non-gridded (surface) resamplings were performed using mri_vol2surf (FreeSurfer).

Many internal operations of fMRIPrep use Nilearn 0.6.2^31^, mostly within the functional processing workflow. For more details of the pipeline, see the section corresponding to workflows in fMRIPrep’s documentation.

**Copyright Waiver**

The above boilerplate text was automatically generated by fMRIPrep with the express intention that users should copy and paste this text into their manuscripts unchanged. It is released under the CC0 license.

The preprocessed fMRI data were further denoised using the Python package Nilearn^31^. The steps included (1) detrending, (2) first-order Butterworth filtering (0.01-0.1Hz), (3) nuisance regression of 36 confounds^32^ (mean WM signal, mean CSF signal, global signal with their temporal derivatives, squares of tissue signals and their temporal derivatives, and 24 head motion parameters^33-34^). Following the recommendation of Lindquist et al.^35^, the nuisance regression was conducted orthogonally to band-pass temporal filtering.

**Tracking modular dynamics of brain network**

In this study, we used a multilayer network framework^37-38^ to track the modular dynamics of functional networks. This framework treats sliding-window functional networks as a continuous-time multilayer network. A generalized Louvain community detection algorithm^39^ was performed to detect dynamic modules of a multilayer network. We used the Genlouvain package to run this algorithm (<https://github.com/GenLouvain/GenLouvain>). This algorithm finds the optimal assignment of community by maximizing the multilayer modularity measure *Q*, which is defined as follows:

$Q(\gamma, \omega) = \frac{1}{2\mu}\sum_{ijsr} [(A_{ijs}-r_{s}\frac{k_{is}k_{js}}{2m_{s}})\delta(M_{is},M_{js}) + \delta(i,j)\omega_{jrs}]\delta(M_{is},M_{jr})$	(1)

For this formula, $\mu$ is the total degree of the multilayer functional network, $A_{ijs}$denotes the connectivity strength between nodes *i* and *j* for time point *s*, $r_{s}$ denotes the topological resolution parameter of time point *s*, $k_{is}$ is the degree of node *i* for time point *s*, $m_{s}$ denotes the total degree of time point *s*, $\delta(x,y)$ denotes the Kronecker delta, $M_{is}$ denotes the community assignment of node *i* for time point *s*, $M_{jr}$ denotes the community assignment of node *j* for time point *s*, $\omega_{jrs}$ denotes the temporal coupling parameter for node *j* between time points *r* and *s*. When completing community assignments for multilayer functional networks, the flexibility of node *i*, $f_{i}$, is defined as follows^37-38,40^:

$f_{i} = \frac{n_{i}}{N}$	(2)

where $N$ indicates the number of possible changes in community, and $n_{i}$ denotes the number of empirical community changes. In the main analysis, we set the topological resolution parameter $\gamma$ to 1, and set the temporal coupling parameter $\omega$ to 1. To solve the pseudo-randomness of the generalized Louvain algorithm, we repeated the community detection 100 times. The mean nodal flexibility over 100 times was used for subsequent analysis.

**Transcriptomic-neuroimaging association analysis**

We used transcriptomic data from the AHBA database^41^, which consists of microarray data in 3702 different brain tissue samples from 6 neurotypical donors (1 female and 5 males, aged from 24 to 57 years). Among the six donors, two have brain tissue samples covering both hemispheres, while four have tissue samples from only the left hemisphere. Thus, we restricted the transcriptomic analysis to the left hemisphere.

We preprocessed the microarray data using the Python package abagen^42-43^. The detained steps included (1) updating the MNI coordinates of all the tissue samples, (2) reannotating microarray probe-to-gene mappings using information provided by Arnatkevic̆iūtė et al.^43^, (3) intensity-based filtering of probes to remove probes that do not exceed background noise in 50% of tissue samples, (4) choosing the probe with the highest differential stability in donors from the probes indexing the same gene, (5) matching tissue samples to regions of functional parcellation we used, (6) normalizing expression data for each sample across genes for each donor using a scaled robust sigmoid normalization function, (7) normalizing expression data for each gene across samples for each donor using a scaled robust sigmoid normalization function, (8) calculating regional expression values by averaging expression values of samples separately for each donor and then averaging across donors. In addition, when a region in the functional parcellation is not assigned a tissue sample in all donors, the region will be assigned the expression values of the tissue sample closest to the centroid of that region. The process eventually yielded expression values of 15633 genes in 158 brain parcels.

We used a partial least squares (PLS) regression to assess the relationship between group differences in flexibility and gene expression. In the PLS regression, gene expression data were treated as predictors of the differences in flexibility. The PLS regression was fitted by the SIMPLS algorithm^45^ using the R package pls^44^. The first PLS component (PLS1), expressed as a linear combination of gene expression data, had the strongest correlation with the differences in flexibility.

**Relation between flexibility and clinical data**

There were 176 patients with PANSS positive scores, 175 patients with PANSS negative scores, and 175 patients with medication information. Based on the formula from Leucht et al.^46^, we converted antipsychotic doses to chlorpromazine (CPZ) equivalents. First, we assessed Pearson correlations between flexibility and clinical data (PANSS positive scores, PANSS negative scores, and CPZ equivalents) at the global and nodal levels. Second, we utilized a support vector regression (SVR) model to determine whether nodal flexibility can predict the PANSS positive and negative scores^47^. We performed leave-one-out cross-validation to assess the accuracy of the model. To avoid information leakage in the model, here we used the original nodal flexibility values (do not perform the NeuroCombat^48^ method to correct for the site effect) and treated site effects as a covariate. In the training set, we used a general linear model (GLM) to remove covariates, including age, sex, mean FD, and site effects, from nodal flexibility values. Similarly, we used a general linear model (GLM) to remove covariates of age and sex from the PANSS scores. The estimated parameters of GLM were further applied to the test set. For the training set, we calculated Pearson correlations between nodal flexibility values and PANSS scores. Nodal flexibility values with a significant correlation (uncorrected p < 0.05) were considered as features. We used the selected features and PANSS scores to fit an SVR model in the training set. Lastly, the trained SVR model was applied to the test set. The model performance was assessed by calculating the Pearson correlation between the predicted and empirical PANSS scores. To determine whether the model performance exceeded the chance level, a total of 10, 000 permutation tests with randomly shuffled PANSS scores were conducted. All results were performed the FDR correction^49^

, with the significance threshold set to corrected p < 0.05. The SVR model was fitted using the Matlab toolbox LIBSVM^50^ (<https://github.com/cjlin1/libsvm>).

**Validation analysis**

We considered multiple confounds that could affect our findings. (1) Since there was no optimal choice of window length in the sliding-window method, we additionally used two different window lengths (40s and 80s) to validate our results. (2) Considering that parameters of a generalized Louvain community detection algorithm may affect our findings, we included three different sets of parameters (($\omega, \gamma$) = (0.75, 1), ($\omega, \gamma$) = (0.75, 1.1), and ($\omega, \gamma$) = (1, 0.9)). (3) We retained the top 15% of the strength of windowed functional connectivity to remove weak or spurious connectivity. Then, we repeated our analysis on the thresholded multilayer functional networks.

**Supplementary tables**

**Table S1: Demographic Characteristics of Participants per site**

|  | HC | | | SZ | | |
| --- | --- | --- | --- | --- | --- | --- |
| Site | N | Age (y) | Sex (F/M) | N | Age (y) | Sex (F/M) |
| COBRE | 54 | 35.6$\pm$10.99 | 14/40 | 31 | 32.7$\pm$12.35 | 6/25 |
| BrainGluSchi | 30 | 28.9$\pm$10.06 | 14/16 | 25 | 32.3$\pm$13.04 | 1/24 |
| UCLA-CNP | 41 | 34.3$\pm$8.59 | 7/34 | 39 | 35.5$\pm$8.91 | 9/30 |
| SWA | 16 | 40.8$\pm$8.67 | 3/13 | 13 | 41.8$\pm$9.10 | 4/9 |
| UTO | 35 | 35.9$\pm$9.57 | 15/20 | 30 | 29.4$\pm$9.83 | 9/21 |
| KUT | 43 | 41.3$\pm$12.46 | 21/22 | 32 | 41.8$\pm$10.73 | 19/13 |
| KTT | 45 | 34.1$\pm$7.91 | 13/32 | 43 | 37.4$\pm$9.98 | 20/23 |
| ds005073\ds003404 | 15 | 35.3$\pm$10.61 | 6/9 | 10 | 40.4$\pm$5.34 | 3/7 |

Note: N denotes the number of participants, Age: Mean$\pm$SD, F: Female, M: male, HC: Healthy controls, SZ: schizophrenia.

**Table S2: Network-level case-control differences**

| Functional network | Cohen’s d value | p-value | Adjusted p-value |
| --- | --- | --- | --- |
| Visual | 0.145 | 0.109 | 0.146 |
| **Somatomotor** | **0.270** | **0.003** | **0.023** |
| Dorsal attention | -0.087 | 0.334 | 0.382 |
| Ventral attention | -0.178 | 0.049 | 0.098 |
| Limbic | -0.185 | 0.040 | 0.098 |
| Frontoparietal | 0.021 | 0.815 | 0.815 |
| Default mode | -0.203 | 0.025 | 0.098 |
| Subcortical | -0.146 | 0.106 | 0.146 |

Note: Adjusted p-value: FDR corrected

**Table S3: Significant nodal-level case-control differences**

| Regions | Cohen’s d value | p-value | Adjusted p-value |
| --- | --- | --- | --- |
| 7Networks_LH_Vis_2 | -0.320 | 0.000433 | 0.0103 |
| 7Networks_LH_SomMot_10 | 0.449 | 0.000001 | 0.0003 |
| 7Networks_LH_SomMot_13 | 0.337 | 0.000209 | 0.0103 |
| 7Networks_LH_SomMot_17 | 0.297 | 0.001078 | 0.0200 |
| 7Networks_LH_SomMot_18 | 0.318 | 0.000458 | 0.0103 |
| 7Networks_LH_SomMot_19 | 0.278 | 0.002217 | 0.0351 |
| 7Networks_LH_SomMot_22 | 0.310 | 0.000648 | 0.0137 |
| 7Networks_LH_SomMot_25 | 0.266 | 0.003323 | 0.0389 |
| 7Networks_LH_SomMot_26 | 0.321 | 0.000416 | 0.0103 |
| 7Networks_LH_SomMot_28 | 0.298 | 0.001014 | 0.0200 |
| 7Networks_LH_Default_Par_2 | -0.265 | 0.003532 | 0.0399 |
| 7Networks_LH_Default_pCunPCC_2 | -0.271 | 0.002827 | 0.0363 |
| 7Networks_LH_Default_pCunPCC_3 | -0.331 | 0.000276 | 0.0103 |
| 7Networks_RH_Vis_3 | 0.277 | 0.002303 | 0.0351 |
| 7Networks_RH_Vis_15 | 0.270 | 0.002872 | 0.0363 |
| 7Networks_RH_Vis_18 | 0.326 | 0.000334 | 0.0103 |
| 7Networks_RH_SomMot_15 | 0.370 | 0.000048 | 0.0048 |
| 7Networks_RH_SomMot_17 | 0.360 | 0.000076 | 0.0048 |
| 7Networks_RH_SomMot_20 | 0.272 | 0.002766 | 0.0363 |
| 7Networks_RH_SomMot_22 | 0.364 | 0.000065 | 0.0048 |
| 7Networks_RH_SomMot_27 | 0.320 | 0.000438 | 0.0103 |
| 7Networks_RH_SomMot_28 | 0.328 | 0.000307 | 0.0103 |
| 7Networks_RH_SalVentAttn_TempOccPar_5 | -0.274 | 0.002529 | 0.0363 |
| 7Networks_RH_SalVentAttn_FrOperIns_5 | -0.292 | 0.001314 | 0.0231 |
| 7Networks_RH_Default_Par_3 | -0.269 | 0.002996 | 0.0364 |
| 7Networks_RH_Default_pCunPCC_1 | -0.324 | 0.000363 | 0.0103 |
| 7Networks_RH_Default_pCunPCC_2 | -0.377 | 0.000035 | 0.0048 |
| 7Networks_RH_Default_pCunPCC_3 | -0.276 | 0.002335 | 0.0351 |

Note: Adjusted p-value: FDR corrected

**Table S4: The top 15 terms relevant to cognitive behaviors or functions from functional decoding analysis based on the NeuroSynth database.**

| Cognitive terms | Pearson’s r | Cognitive terms | Pearson’s r |
| --- | --- | --- | --- |
| motor | 0.2456 | episodic | 0.1387 |
| primary motor | 0.1892 | autobiographical | 0.1279 |
| sensorimotor | 0.1630 | retrieval | 0.1019 |
| movements | 0.1533 | episodic memory | 0.0983 |
| premotor | 0.1431 | memory | 0.0933 |
| hand | 0.1413 | autobiographical memory | 0.0899 |
| somatosensory | 0.1163 | construction | 0.0853 |
| finger | 0.1154 | memories | 0.0803 |
| supplementary motor | 0.1148 | events | 0.0794 |
| foot | 0.0938 | memory retrieval | 0.0754 |
| primary sensorimotor | 0.0898 | remembering | 0.0721 |
| motor imagery | 0.0810 | navigation | 0.0690 |
| coordination | 0.0806 | past | 0.0656 |
| motor premotor | 0.0799 | future | 0.0631 |
| primary somatosensory | 0.2456 | semantic memory | 0.0602 |

Note: The first two columns show functional decoding results for regions with higher flexibility, and the last two columns show functional decoding results for regions with lower flexibility.

**Table S5-S6: Please check sheets in “Supplementary table S5-S6.xlsx”**

**Supplementary figures**

**Figure S1.** Spearman correlations between the PLS1 weights of all genes and the differential gene expression of all genes reported by Gandal et al. and Fromer er al.. All p values were performed the FDR correction, with the significance threshold set to corrected p < 0.05. SZ: schizophrenia, BD: bipolar disorder, ASD: autism spectrum disorder, MDD: major depression disorder, AAD: alcohol abuse disorder, IBD: inflammatory bowel disease.

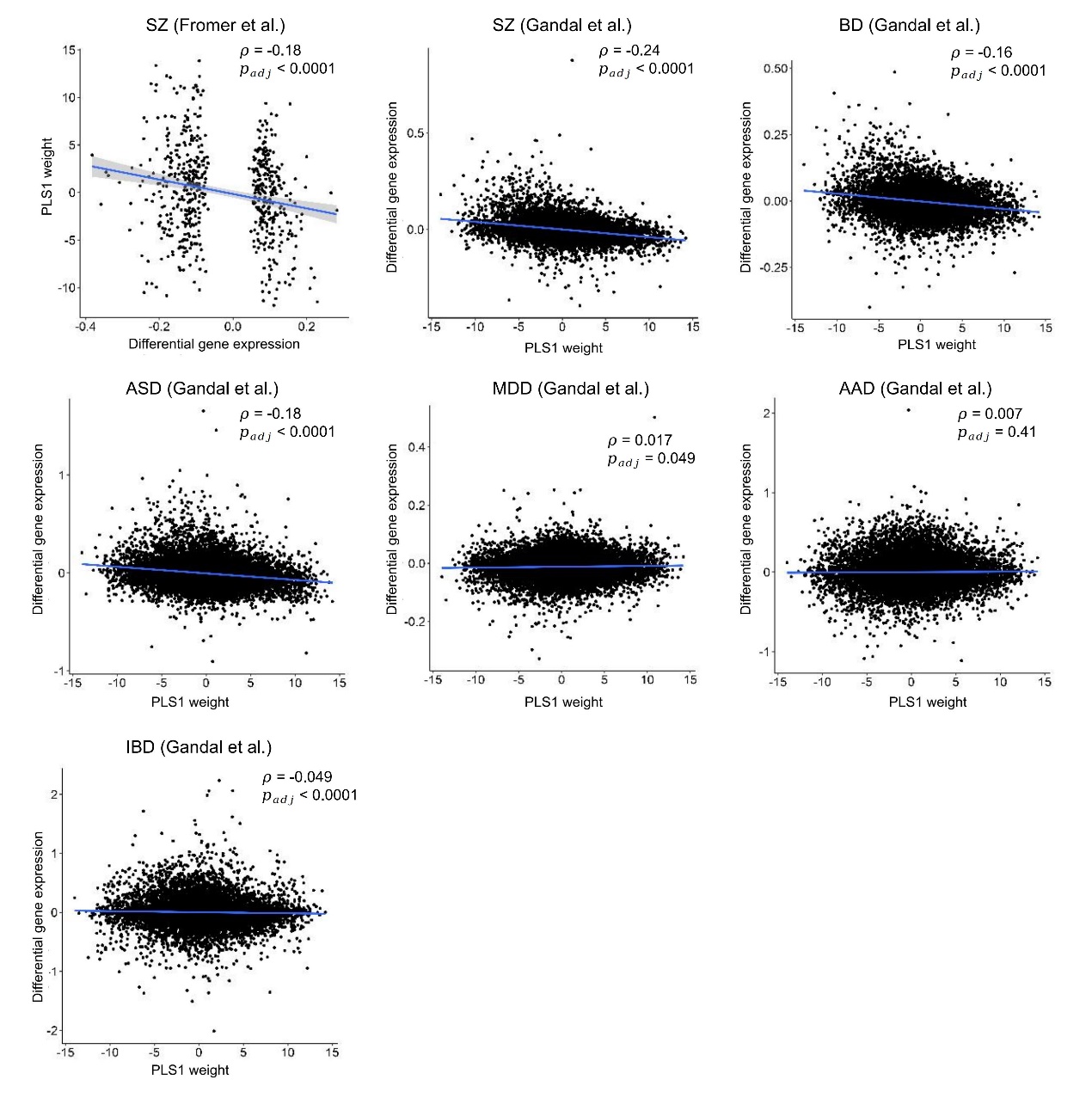

**Figure S2.** Correlations between empirical PANSS positive scores and predicted PANSS positive scores, as well as between empirical PANSS negative scores and predicted PANSS negative scores. P values were corrected by the FDR method.

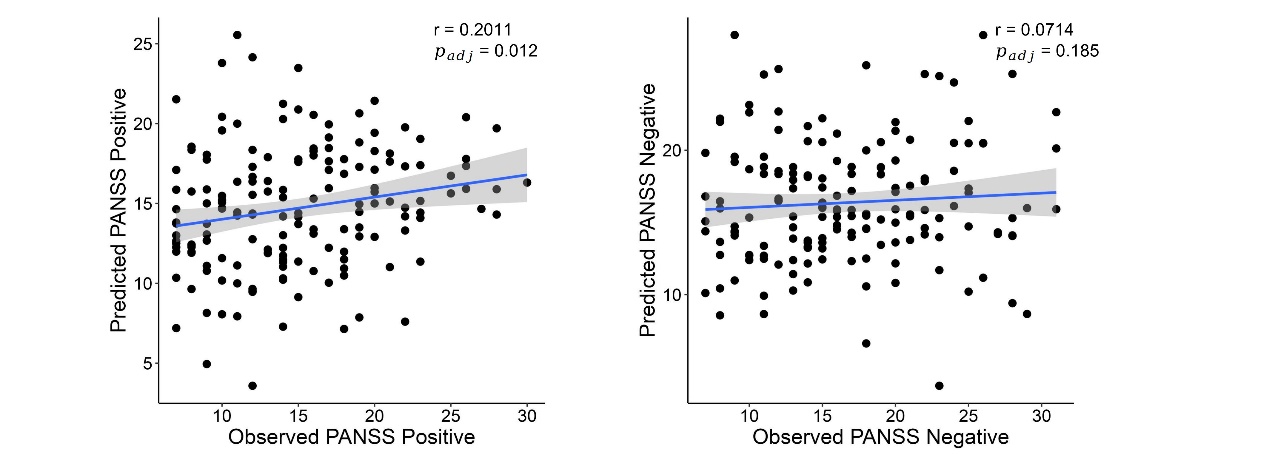

**Figure S3-S8.** Results of validation analysis. We repeated the analysis of case-control comparisons under multiple confounds, including different sliding-window length (40s and 80s), different parameter selections of the generalized Louvain algorithm (($\omega, \gamma$) = (0.75, 1), ($\omega, \gamma$) = (0.75, 1.1), and ($\omega, \gamma$) = (1, 0.9)), and using thresholded windowed functional connectivity. For each validation analysis, (a). SZ-HC differences in mean flexibility in the validation analysis, (b). The cohen’s d map of case-control comparisons in the validation analysis, (c). Significant SZ-HC differences in flexibility at the node level in the validation analysis (FDR-corrected p < 0.05). (d). The correlation between SZ-HC differences in the main analysis and SZ-HC differences in the validation analysis.

**Figure S3: Window length = 40s**

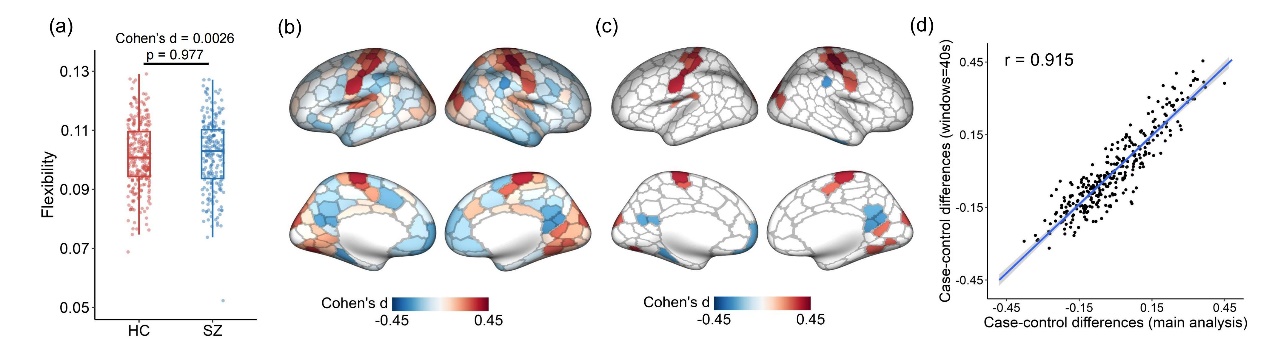

**Figure S4: Window length = 80s**

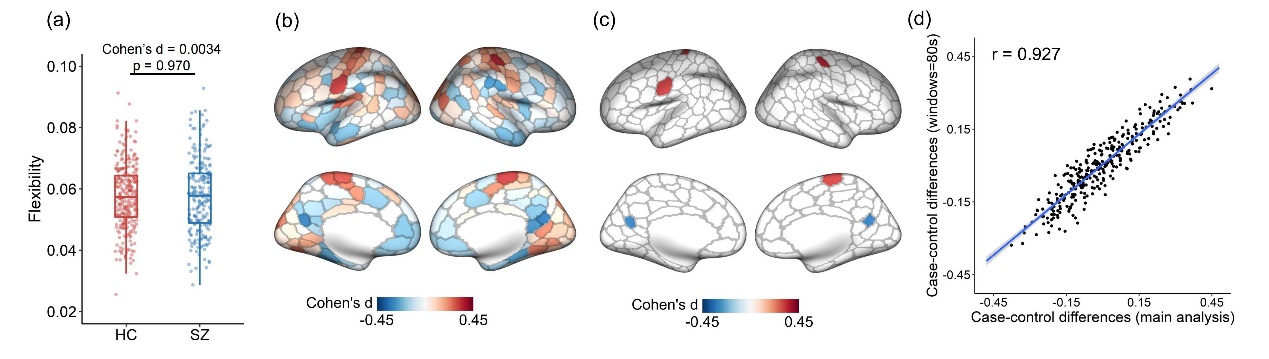

**Figure S5:** ($\omega, \gamma$) = (0.75, 1)

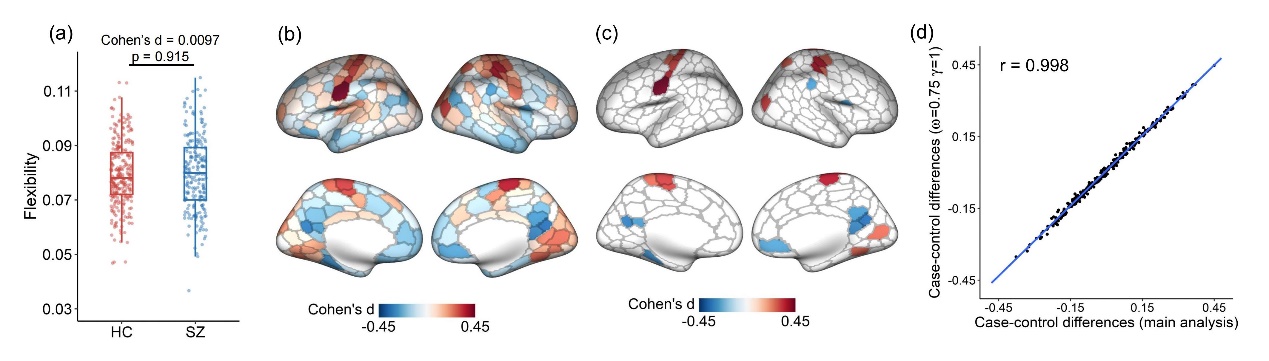

**Figure S6:** ($\omega, \gamma$) = (0.75, 1.1)

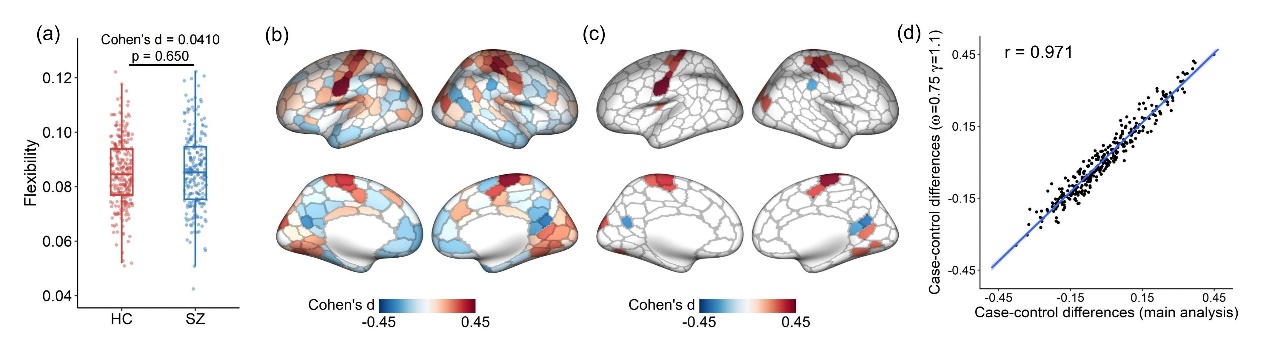

**Figure S7:** ($\omega, \gamma$) = (1, 0.9)

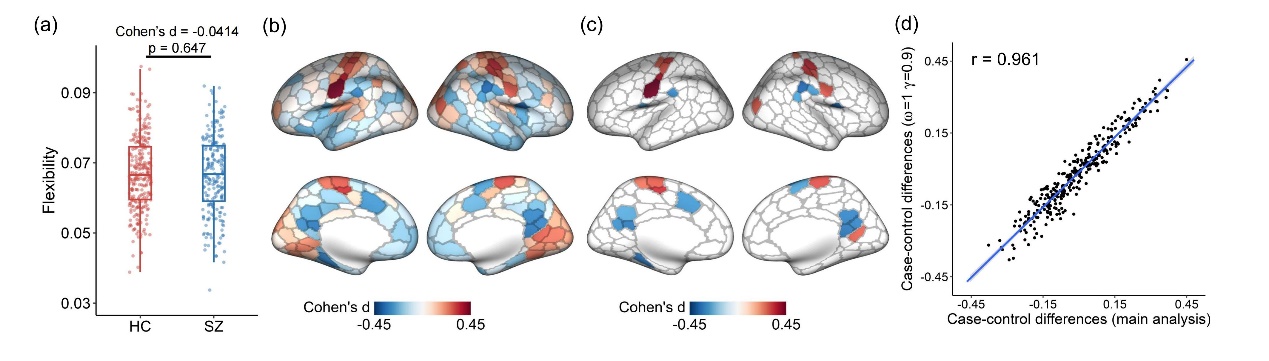

**Figure S8: Thresholded windowed functional connectivity**

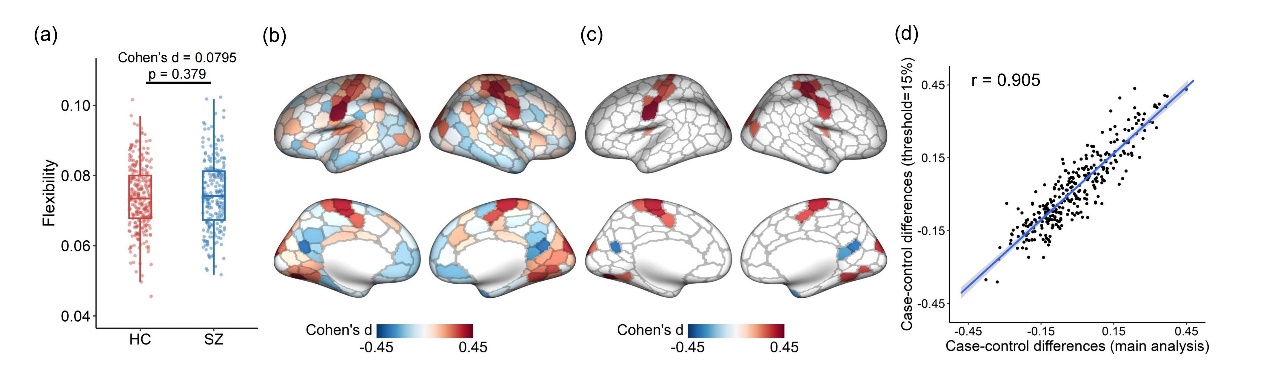
